# Seascape genomics reveals population structure and local adaptation in a widespread coral reef snail, *Coralliophila violacea* (Kiener, 1836)

**DOI:** 10.1101/2021.06.15.448144

**Authors:** Sara E. Simmonds, Samantha H. Cheng, Allison L. Fritts-Penniman, Gusti Ngurah Mahardika, Paul H. Barber

## Abstract

Local adaptation to different environments may reinforce neutral evolutionary divergence, especially in populations in the periphery of a species’ geographic range. Seascape genomics (high-throughput genomics coupled with ocean climate databases) facilitates the exploration of neutral and adaptive variation in concert, developing a clearer picture of processes driving local adaptation in marine populations. This study used a seascape genomics approach to test the relative roles of neutral and adaptive processes shaping population divergence of a widespread coral reef snail, *Coralliophila violacea*. We collected *C. violacea* from colonies of their coral host (*Porites* spp.) at ten locations spanning a large portion of their geographic range. We used RAD-seq to investigate possible local adaptation via genetic-environmental associations with five ocean climate variables. Four genetic partitions were concordant with regions previously observed in mtDNA (Indian Ocean, Coral Triangle, and Hawaiian Islands), with the addition of Vietnam and varying isolation and admixture levels. We identified outlier loci (FDR = 0.10, *N* = 72) among individual localities and between regions (FDR = 0.10, *N* = 34), suggesting that some loci are putatively under divergent selection. Association analyses showed that the two strongest drivers of local adaptation were the annual range and mean of sea surface temperature. Populations that experience lower sea surface temperatures at the periphery of *C. violacea’s* geographic range drive these associations. Our results show that local adaptation to different environments likely reinforces neutral divergence, especially in peripheral populations.

## INTRODUCTION

Incomplete or ephemeral dispersal barriers, combined with potentially large population sizes and extensive geographic ranges of marine taxa, suggest that natural selection plays a crucial role in generating biodiversity in the ocean (Crandall et al., 2012). Authors are increasingly advocating for speciation models that combine both neutral and adaptive processes (Bowen et al., 2013; Horne, 2014) because natural selection limits realized connectivity between populations (Burgess et al., 2012). Local adaptation to climate and habitat can reinforce neutral divergence patterns driven by gene flow, genetic drift, or mutations (Gavrilets, 2003). In these models, even species with high fecundity and long dispersal distances can have low effective gene flow if selection favors local progeny over those recruiting from different environments.

Empirical evidence indicates that selection plays a more significant role in the diversification of marine taxa than previously thought. For example, several studies show environmental and habitat heterogeneity (Longo & Bernardi, 2015; Rocha & Bowen, 2008; Teske et al., 2019), or competition among species (Bowen et al., 2013; Briggs, 1992), drive diversification or reinforce nascent allopatric divergence. Similarly, recent work also suggests that divergent selection between habitats or hosts contributes to adaptive variation and speciation (Cheng, 2015; Meyer et al., 2005; Reid et al., 2006; Simmonds et al., 2020; Tornabene et al., 2015).

Natural selection can be particularly influential in peripheral areas of species’ ranges where environmental conditions may be at or near the limits of a species’ physiological threshold (J. D. DiBattista et al., 2016; Gaither et al., 2010; Johannesson & André, 2006; Kawecki, 2008), creating different selective pressures.

Studies using high-throughput genomic tools have found signals of natural selection driving genetic divergence in peripheral marine populations (Ackiss et al., 2018; Gaither et al., 2015; Saenz-Agudelo et al., 2015). Authors hypothesized various mechanisms driving divergence, including different ecological (e.g., habitats) and environmental factors (e.g., sea surface temperature, salinity, turbidity), but also seasonal fluctuations or environmental heterogeneity (e.g., oceanography) at smaller scales that indicate selective processes at play. As such genomic sequencing, combined with global, high-resolution marine environmental databases (Sbrocco & Barber, 2013), meaning we can now directly assess the role of natural selection driving or reinforcing diversification in the ocean.

In recent decades, geneticists have documented phylogeographic structure in a wide diversity of marine taxa (Bowen et al., 2014; Kelly & Palumbi, 2010), advancing allopatry as a primary speciation model in marine ecosystems, particularly among studies of Indo-Pacific biodiversity (Barber et al., 2011; Bowen et al., 2013; Carpenter et al., 2010; Gaither et al., 2010). Authors often report one or more of the following phylogeographic patterns: 1) population divergence between Indian and Pacific Ocean basins; 2) population differentiation within the Coral Triangle, the global epicenter of marine biodiversity; or 3) population differentiation on the periphery of the Pacific (e.g., Hawai’i, Marquesas) or Indian Oceans (e.g., Red Sea).

The divergence between the Indian and Pacific Oceans is typically ascribed to the Sunda and Sahul continental shelves’ exposure when Plio-Pleistocene glaciations lowered sea levels by 115–130m (Voris, 2000). These intermittent landmass barriers constricted waterways of the Indonesian and Philippine Archipelago, reducing gene flow between the Indian and Pacific Oceans in a wide diversity of marine taxa, including seahorses (Lourie et al., 2005); soldierfish (Craig et al., 2007); anemonefish (Dohna et al., 2015; Timm & Kochzius, 2008); damselfish (Drew & Barber, 2009; Liu et al., 2014; Raynal et al., 2014; Liu et al. 2019); groupers (Gaither et al., 2011); fusiliers (Ackiss et al., 2013); limpets (Kirkendale & Meyer, 2004); snails (Crandall et al., 2008a; Reid et al., 2006; Simmonds et al., 2018); giant clams (DeBoer et al., 2014; Nuryanto & Kochzius, 2009) and seastars (Kochzius et al., 2009). Further studies suggest that modern oceanographic features such as the Halmahera Eddy can limit gene flow within the Coral Triangle by constraining larval exchange, a hypothesis supported by both phylogeographic studies (Ackiss et al., 2013; Barber et al., 2011; DeBoer et al., 2008) and biophysical connectivity models (Kool et al., 2011; Treml et al., 2015).

Although multiple processes contribute to isolation and divergence in the sea (Barber & Meyer 2014), purely allopatric models of marine speciation still face challenges. For example, while Plio-Pleistocene sea levels did constrict Indonesian and Philippine waterways (Voris, 2000), the major pathways for the Indonesian Throughflow, the Makassar Strait, Maluku, and Banda Seas, are deeper than 3,000 m. Thus, even at the lowest sea levels, these dispersal pathways between Pacific and Indian Oceans remained open, requiring authors to invoke auxiliary processes such as increased upwelling of cold water as a mechanism to limit dispersal (Fleminger, 1986). Moreover, periods of isolation were punctuated by tens of thousands of years of oceanic conditions similar to today’s, during which gene flow and population expansion would have occurred, weakening the signal of historical isolation (Crandall et al., 2008b).

Similarly, ocean currents in the region vary, both seasonally (Shinoda et al., 2012) and across epochs (Kuhnt et al., 2004), such that oceanographic features may promote isolation during some periods only to be reversed in others.

The coral-eating snail *Coralliophila violacea* (Kiener, 1836) is widespread on coral reefs across the tropical Indo-Pacific, from the Red Sea to the Eastern Pacific (Demond, 1957), and may be subject to both neutral and selective processes for divergence. Previous work on *C. violacea* demonstrated genetic differentiation of snails living on different coral host lineages, despite ongoing gene flow (Simmonds et al., 2020; Simmonds et al., 2018). One lineage was broadly distributed but exhibited phylogeographic structure in the core of its geographic range (i.e., across the Sunda Shelf) and peripheral populations (i.e., Hawai i). This structure could be attributed to allopatric processes, such as isolation during low sea-level stands or isolation by distance. However, given that environmental conditions vary greatly, it is possible that variation in ocean climate across the range of *C. violacea* could reinforce population divergence via natural selection.

This study combined genome-wide surveys of genetic variation in *C. violacea* with data from marine environmental databases to test the relative roles of neutral and adaptive processes shaping population divergence. Specifically, we test for divergence across known phylogeographic provinces within the Coral Triangle and divergence among peripheral populations in the Indian and Pacific Oceans and the South China Sea. We then compare neutral and non-neutral variation patterns to geography and environmental variables to assess their relative roles in shaping population divergence.

## MATERIALS AND METHODS

### Sample collection

To test for patterns of divergence related to geography and local adaptation in *C. violacea*, we collected snails on snorkel or scuba at ten locations spanning a large portion of the snail’s geographic range (Table 1, Fig. 1). Sampled populations included both sides of the Sunda Shelf, an area where phylogeographic structure is commonly observed, as well as Hawai i, the edge of *C. violacea’s* geographic range and an area known to exhibit peripheral isolation (Gaither et al., 2011b; Hodge et al., 2014). In addition, we only used snails collected from *P. lobata* and *P. compressa* coral colonies (clades 2 and 3, the “green group” from Simmonds et al. 2018) that were previously DNA barcoded using RAD-seq data. This selectivity was necessary because *Porites* corals are notoriously challenging to identify *in situ* due to their morphological plasticity and small corallites (Forsman et al., 2015) and because genetically similar colonies can have vastly different morphologies and *vice versa* (Forsman et al., 2009). In total, we collected snails from 1–3 colonies at each location for a total of 71 snails from 32 coral colonies. A portion of each snail’s foot tissue was preserved in 95% ethanol and stored at room temperature for DNA analysis.

**Table 1.** Sampling locations for *Coralliophila violacea* collected from coral hosts *Porites lobata* and *P. compressa* from *Porites* lineage 1, corresponding environmental variables from MARSPEC database for each location used in Bayenv2 analysis. Coordinates are in decimal degrees. Location numbers correspond to those in Figure 1. Regions were used for AMOVA analyses. Sea surface salinity (SSS) in psu (practical salinity units). Sea surface (SST) in °C. *N* = no. of individuals.

| LOCATION | COUNTRY | REGION | LATITUDE | LONGITUDE | WARMEST MO. |  | SST | SST RANGE | SSS | SSS RANGE |
| --- | --- | --- | --- | --- | --- | --- | --- | --- | --- | --- |
|  |  |  |  |  | SST |  |  |  |  |  |
| 1. Vavvaru | Maldives | Indian Ocean | 5.419 | 73.358 | 29.42 |  | 28.37 | 1.76 | 34.93 | 1.64 |
| 2. Pulau Weh | Indonesia | Indian Ocean | 5.887 | 95.348 | 29.39 |  | 28.59 | 1.40 | 33.29 | 1.43 |
| 3. Hòn Mun | Vietnam | Vietnam | 12.170 | 109.308 | 28.20 |  | 26.45 | 4.22 | 33.34 | 0.77 |
| 4. Dumaguete | Philippines | Coral Triangle | 9.332 | 123.312 | 28.54 |  | 27.61 | 3.16 | 33.96 | 0.92 |
| 5. Bunaken | Indonesia | Coral Triangle | 1.612 | 124.783 | 28.74 |  | 28.34 | 1.05 | 33.95 | 0.79 |
| 6. Lembeh | Indonesia | Coral Triangle | 1.479 | 125.251 | 28.81 |  | 27.87 | 2.94 | 33.95 | 0.85 |
| 7. Pulau Mengyatan | Indonesia | Coral Triangle | -8.557 | 119.685 | 28.47 |  | 27.51 | 1.95 | 34.09 | 0.68 |
| 8. Nusa Penida | Indonesia | Coral Triangle | -8.675 | 115.513 | 28.81 |  | 27.60 | 2.12 | 33.89 | 1.19 |
| 9. Manokwari | Indonesia | Coral Triangle | -0.888 | 134.085 | 29.16 |  | 28.90 | 0.78 | 34.09 | 1.03 |
| 10. Ka'a'awa | USA | Hawai'i | 21.584 | -157.887 | 26.43 |  | 24.99 | 2.81 | 34.92 | 0.21 |

**Figure 1.**
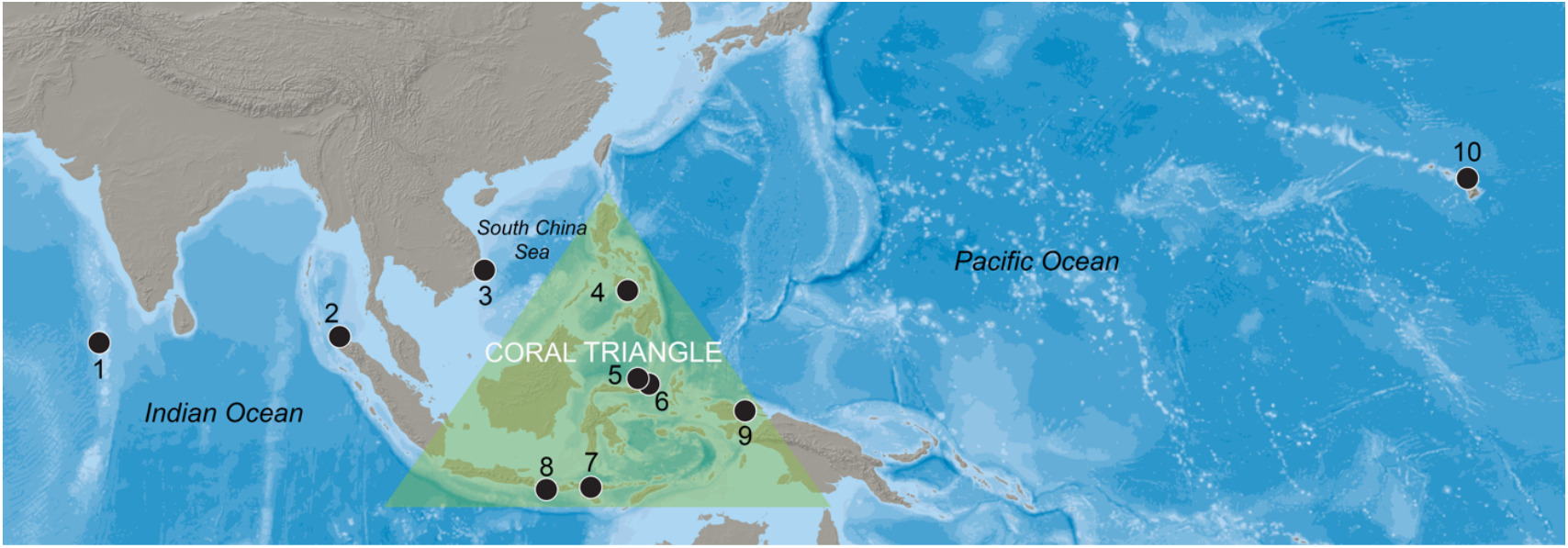
Map of *Coralliophila violacea* collection locations across the Indo-Pacific region. Sampled populations include 1. Vavvaru. 2. Pulau Weh 3. Hòn Mun 4. Dumaguete 5. Bunaken 6. Lembeh 7 Pulau Mengyatan 8. Nusa Penida 9. Manokwari 10. Ka’a’awa. Raster map made with Natural Earth.

### Creation of RAD-seq libraries

We extracted DNA from 20 mg of foot tissue using a Qiagen DNeasy Blood and Tissue Kit (Qiagen). We followed the manufacturer’s instructions, except in the last step, when we eluted DNA with 100 µl of molecular grade H_2_O rather than AE buffer. We estimated initial DNA concentrations using a NanoDrop and visualized DNA quality on a 1% agarose gel stained with SYBR Green. We used only high-quality DNA with a bright high molecular weight band and minimal smearing. We dried DNA extractions using a SpeedVac on medium heat and reconstituted using molecular grade H_2_O to a final uniform 250 ng/µl DNA concentration.

We prepared 2b-RAD libraries following the protocol of Wang et al. (Wang et al., 2012). AlfI restriction enzymes digest reduced representation (1/16^th^) libraries labeled with individual barcodes and subjected to 18–20 PCR amplification cycles. We electrophoresed products on a 2% agarose gel in a 1× TBE buffer and run at 150 V for 90 minutes. Target bands (165 bp) were visualized with SYBR SAFE dye and excised from the gel. The excised band was then purified using a QIAquick gel purification kit (Qiagen). For a final cleaning step, we used Ampure XP beads (Beckman-Coulter). The QB3 Vincent J. Coates Genomics Sequencing Laboratory at UC Berkeley conducted quality checks (qPCR, BioAnalyzer) and sequencing of the resulting libraries, multiplexing 10–20 snails per lane in 5 lanes of a 50 bp single-end run on the Illumina HiSeq 2000 platform.

### RAD-seq data processing

To prepare raw sequence data for SNP identification, we truncated all raw reads to the insert size (36 bp), filtered for quality (PHRED scores >20), and discarded empty constructs. We used custom scripts to process the data, available on GitHub https://github.com/z0on/2bRAD_denovo. First, unique tag sequences (minimum sequencing depth 5×) were counted, the number in reverse-complement orientation recorded and merged into one table. All sequences were then clustered in CD-HIT using a 91% similarity threshold. Next, we defined the reference as the most abundant sequence in the cluster. We then filtered a locus-annotated table from the previous two steps, excluding reads below 5× depth or exhibiting strand bias. The resulting clustered sequences’ orientation was then flipped to match the most abundant tag in a cluster.

We applied mild allele filters (10× total depth, allele bias, and strand bias), with the requirement that alleles appear in at least two individuals. We then applied locus filters, allowing a maximum of 50% heterozygotes at a locus, no more than two alleles, genotyped in 30% of samples, and polymorphic. The final set of SNPs was then thinned to one per tag (the one with the highest minor allele frequency) for STRUCTURE analysis and gene-environment association tests to remove linked loci that might be in linkage disequilibrium.

### Individual sample filtering steps

From the first 71 individuals, we filtered out those with low genotyping rates (*N* = 5), indicating poor DNA quality by taking the log_10_ of the number of sites genotyped per individual and removing individuals ≥ 2 standard deviations (SD) of the mean. We also removed individuals (*N* = 4) with high homozygosity (+/– 2 SD of the mean *F* inbreeding coefficient), indicating potential contamination. We used the remaining 63 individuals in all analyses. The final data file was in VCF format, and we converted it to other data formats using PGDSpider v2.1.0.1 (Lischer & Excoffier, 2012).

### Genetic diversity

We calculated basic population genetic summary statistics using all filtered loci for each location with ARLEQUIN v3.5 (Excoffier & Lischer, 2010). We estimated the number and frequency of polymorphic loci for each site, observed and expected heterozygosity, and genetic diversity (averaged across all loci).

### *F*_ST_ outlier loci test

We tested loci potentially under natural selection by identifying outlier *F*_ST_ values using BayeScan v2.1 (Foll & Gaggiotti, 2008). We ran BayeScan with the data structured in two ways to look for outliers at different spatial scales, 1) comparing individual localities and 2) comparing localities grouped into regions (Table 1). We ran the data with a burn-in of 50,000, a thinning interval 10, a sample size 5,000, 100,000 iterations, and 20 pilot runs of 5,000, each examining two different false discovery rates (FDR = 0.10, 0.05).

### Population genetic structure

We used the Bayesian model-based clustering method STRUCTURE (Pritchard et al., 2000) to infer the population genetic structure and individual admixture proportions. We examined three datasets consisting of 63 individuals: 1) full RAD-seq dataset using all loci; 2) using only neutral loci as determined by BayeScan; and 3) outlier loci with a false discovery rate of 10% (FDR = 0.10, BayeScan). We ran STRUCTURE with a burn-in period of 20,000 followed by 50,000 MCMC replicates for *K* = 1–10, and ten runs for each *K*. We used the admixture model, with allele frequencies correlated among populations. We selected the optimal value of *K* using the Δ*K* statistic (Evanno et al., 2005) and summarized the results graphically using the program CLUMPAK v1.1 (Kopelman et al., 2015). To test for the significance of genetic partitions identified by STRUCTURE, we ran analyses of molecular variance (AMOVA), both with and without regional groupings (Table 1). We determined significance by 100,000 random replicates in ARLEQUIN.

### Genetic-environment association tests

Because ocean climate variables that impact species distributions (Briggs, 2006) may also structure population genetics (Sanford & Kelly, 2011), we examined differences in allele frequencies associated with environmental variables using a Bayesian framework (Bayenv 2.0; (Coop et al., 2010; Günther & Coop, 2013) that accounts for demographic history. First, we obtained ocean environmental variables from the MARSPEC database (Sbrocco & Barber, 2013). Because of strong correlations among environmental variables, we selected only five in the MARSPEC database for analysis: 1) temperature of the warmest ice-free month (biogeo15), 2) mean annual sea surface temperature (annual sea surface temperature; biogeo13), 3) annual range in sea surface temperature (biogeo16), 4) mean annual sea surface salinity (biogeo08), and 5) annual range in sea surface salinity (biogeo11). Temperature variables are particularly important to this system because coral reefs in locations with high sea surface temperatures are most likely to be impacted by coral bleaching events (Hoegh-Guldberg, 1999). Thus, coral mortality from bleaching could directly impact *C. violacea* living on coral hosts and indirectly by limiting the number and species of hosts available to *C. violacea* in affected reefs.

To create maps of each of the five environmental variables across the study region, we projected the annual mean and range of sea surface salinity (SSS) and temperature (SST), plus the mean warmest monthly temperature at ∼5 km resolution onto an equidistant cylindrical world using the ‘*raster’* package in R. Then, we extracted the five climate variable data to each point location and divided it by a scaling factor of 100 using custom R scripts. Next, we estimated a covariance matrix with standardized environmental variables for each sampling location, as suggested by the authors of Bayenv2 (Coop et al., 2010; Günther & Coop, 2013), and 3,186 loci that were polymorphic among all sites, for 100,000 iterations, outputting the results every 500 iterations. Finally, we used the last printed covariance matrix for all further analyses. To estimate the Bayes Factor (BF) for each SNP with each ocean climate variable, we ran Bayenv2 for 100,000 Markov chain Monte Carlo (MCMC) iterations. For each ocean climate variable, SNPs with log_10_ BF > 1 were considered to give substantial-to-strong support for environmentally-associated loci, based on criteria from (Litton & Jefferys, 1984).

### Candidate gene annotation

To annotate the function of genes linked to outlier loci and environmentally-associated loci, we aligned sequences containing outlier loci to nucleotides (nr/nt) on the NCBI website using the BLASTN algorithm at two different taxonomic levels, 1) Mollusca (taxid:6447) and 2) Lophotrochozoa (taxid:1206795). We adjusted parameters (word size 7, expected threshold 10, no mask for lookup table, no low complexity filter) to accommodate short-read sequences. We only examined hits with a high percent query coverage (>85%) and used NCBI to identify and annotate any associated genes.

## RESULTS

The average number of unique reads per individual was 6.6 million at minimum 5× depth after filtering for quality and removing empty constructs. We sequenced and genotyped 46,148 high-quality RAD-seq loci with ≥25× coverage in 71 individuals at ten locations. First, we filtered the dataset for 30% maximum missing data per locus, leaving 7,862 loci, and then thinned to one SNP per loci to remove any physically linked SNPs for STRUCTURE and *F*_ST_ analyses, leaving 3,188 SNPs. Next, we removed nine individuals with potential contamination issues (inbreeding coefficient ≥ +2SD from the mean) or low DNA quality (missing data ≥ +1SD from the mean), leaving 63 individuals.

### Genetic diversity

Across all locations, a mean of 47% of loci were polymorphic. Hòn Mun in Vietnam had the highest frequency of polymorphic loci (81%) and the most sequenced individuals (Table 2). Observed heterozygosity (H_O_ = 0.196 – 0.570) and expected heterozygosity (H_E_ = 0.225 – 0.586) varied across locations, but H_O_ was consistently lower than the H_E_ at every location (Table 2). Mean genetic diversity over all loci across sites was 0.122, and genetic diversity varied across locations (0.089 – 0.162; Table 2). Genetic diversity was highest in Dumaguete (0.162) and lowest in Bunaken (0.095) (Table 2).

**Table 2.** Standard diversity indices and summary statistics for all 3,188 loci obtained from *Coralliophila violacea. N* = no. of individuals, no. of polymorphic loci per location, frequency of polymorphic loci, observed heterozygosity (H_O_), expected heterozygosity (H_E_), and gene diversity over all loci.

| Location | $N$ | No. poly. loci | Freq. poly. loci | $H_O$ | $H_E$ | Gene diversity |
| --- | --- | --- | --- | --- | --- | --- |
| 1. Vavvaru | 4 | 1293 | 41% | 0.345 | 0.449 | 0.104 |
| 2. Pulau Weh | 3 | 1051 | 33% | 0.459 | 0.544 | 0.103 |
| 3. Hòn Mun | 19 | 2598 | 81% | 0.196 | 0.225 | 0.159 |
| 4. Dumaguete | 2 | 1022 | 32% | 0.570 | 0.586 | 0.162 |
| 5. Bunaken | 7 | 1577 | 49% | 0.281 | 0.342 | 0.095 |
| 6. Lembeh | 6 | 1515 | 48% | 0.301 | 0.341 | 0.114 |
| 7. Pulau Mengyatan | 4 | 1143 | 36% | 0.360 | 0.463 | 0.089 |
| 8. Nusa Penida | 9 | 1881 | 59% | 0.271 | 0.296 | 0.115 |
| 9. Manokwari | 5 | 1680 | 53% | 0.274 | 0.357 | 0.149 |
| 10. Ka'a'awa | 4 | 1229 | 39% | 0.353 | 0.397 | 0.132 |

### Gene flow and genetic structure

Results from STRUCTURE using the full dataset of 3,188 loci or just 3,116 putatively neutral loci confirmed that *K* = 4 was the best-supported number of population partitions (Evanno et al. 2005). The four genetic partitions corresponded to recognized distinct biogeographic regions: a) Indian Ocean (1. Vavvaru, 2. Pulau Weh); b) Vietnam (3. Hòn Mun); c) the Coral Triangle (4. Dumaguete, 5. Bunaken, 6. Lembeh, 7. Pulau Mengyatan, 8. Nusa Penida and 9. Manokwari); and d) Hawai i (10. Ka’a’awa) (Fig. 2). Levels of admixture among these regions were variable. Hawai i and Vietnam showed minimal admixture from other locations. In contrast, sites in the Indian Ocean had the highest levels of admixture, with numerous individuals exhibiting a mix of alleles from the Indian Ocean and populations to the east. Specifically, Pulau Weh exhibited substantial admixture with the Coral Triangle, and Vavvaru in the Maldives had admixture from both Vietnam and the Coral Triangle. Levels of admixture varied across sites in the Coral Triangle. Pulau Mengyatan and Manokwari had the highest levels, while Dumaguete, Bunaken, Lembeh, and Nusa Penida had the lowest; admixture in these populations was mainly from the Indian Ocean and Vietnam. Results were virtually identical whether we used all loci or neutral loci.

**Figure 2.**
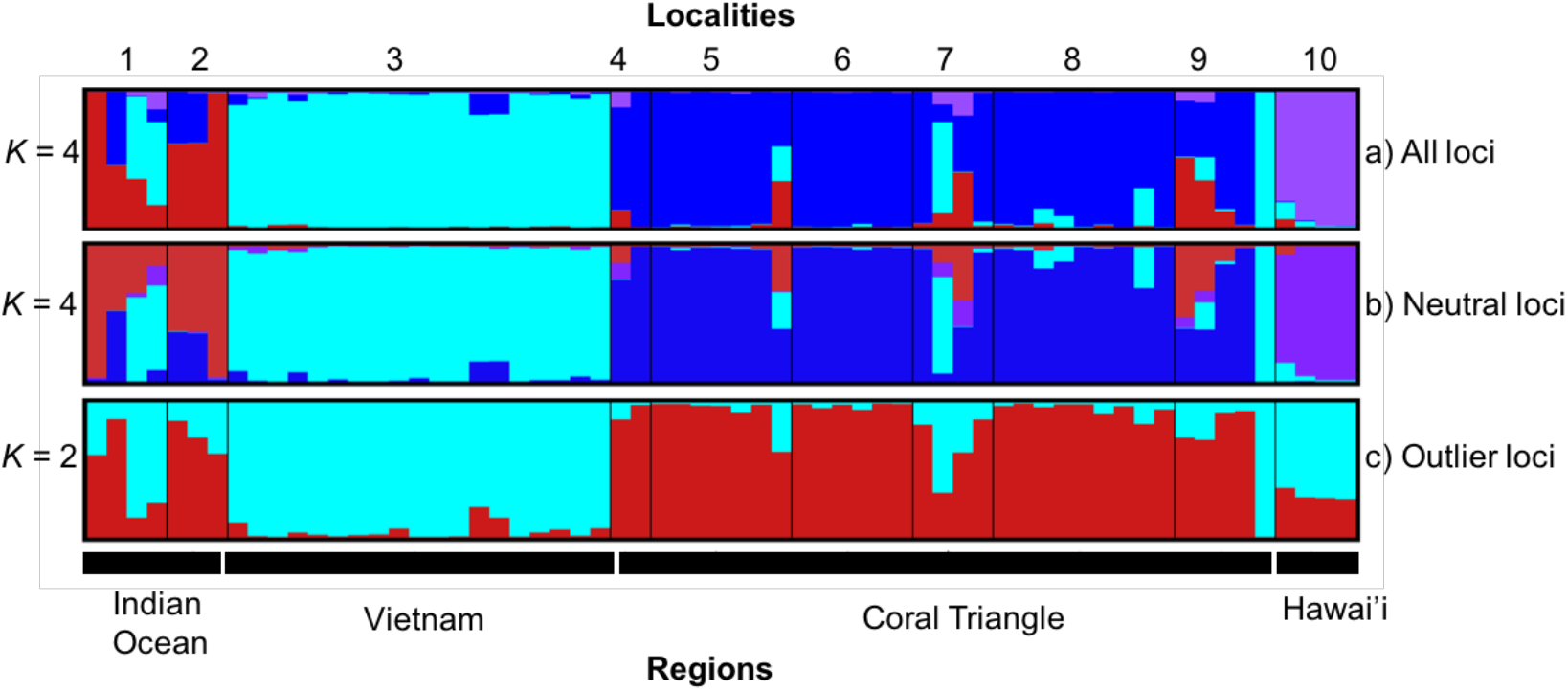
Patterns of genetic structure in *Coralliophila violacea* in the Indo-Pacific region. CLUMPAK-averaged STRUCTURE plots for ten independent runs, clustered and average using CLUMPAK. Each bar represents an individual, and the color is the proportion of assignment to each genetic partition. Locations plotted from left to right and numbered above, and black bars indicate regions as in Table 3.1 **a)** all 3,188 loci at *K* = 4, **b)** 3,116 neutral loci at *K* = 4, **c)** 72 outlier loci at *K* = 2.

Consistent with STRUCTURE results, AMOVA analyses showed significant population structure across all loci (overall *F*_ST_ = 0.071, p >0.001). Hòn Mun in Vietnam was isolated from all other locations (pairwise *F*_ST_ = 0.063 – 0.147; Table 3) except Pulau Mengyatan in southern Indonesia. Ka’a’awa in Hawai i was also strongly divergent from four other locations (pairwise *F*_ST_ = 0.142 – 0.235; Table 3). Similarly, Pulau Weh in the Indian Ocean was different from all locations (pairwise *F*_ST_ = 0.021 – 0.168; Table 3) except for Vavvaru in the Maldives, Dumaguete in the Philippines, and Ka’a’awa in Hawai i. AMOVA analyses grouping sites into four biogeographic regions (Indian Ocean, South China Sea, Coral Triangle, Hawai i; Table 1, Fig. 3) increased the overall *F*_ST_ to 0.098 (p >0.001), and the *F*_CT_ was 0.109 (p >0.001).

**Table 3.** Genetic distances of pairwise populations of *Coralliophila violacea* for all 3,188 SNPs. Regions follow labeling in Table 1.1. Significant pairwise *F*_ST_ values at *p* >0.01 (α=0.05, corrected for multiple tests using the B-Y method, (Narum, 2006) are shaded.

| Location | Indian Ocean |  | Vietnam | Coral Triangle |  |  |  |  |  | Hawai'i |
| --- | --- | --- | --- | --- | --- | --- | --- | --- | --- | --- |
|  | 1 | 2 | 3 | 4 | 5 | 6 | 7 | 8 | 9 | 10 |
| 1. Vavvaru | 0 |  |  |  |  |  |  |  |  |  |
| 2. Pulau Weh | -0.026 | 0 |  |  |  |  |  |  |  |  |
| 3. Hòn Mun | 0.014 | **0.034** | 0 |  |  |  |  |  |  |  |
| 4. Dumaguete | 0.063 | 0.021 | **0.122** | 0 |  |  |  |  |  |  |
| 5. Bunaken | **0.068** | **0.072** | **0.067** | 0.011 | 0 |  |  |  |  |  |
| 6. Lembah | **0.085** | **0.077** | **0.109** | 0.008 | 0.007 | 0 |  |  |  |  |
| 7. Pulau Mengyatan | 0.022 | 0.033 | -0.004 | -0.025 | -0.005 | 0.002 | 0 |  |  |  |
| 8. Nusa Penida | **0.059** | **0.052** | **0.098** | 0.013 | 0.000 | 0.011 | -0.020 | 0 |  |  |
| 9. Manokwari | -0.005 | -0.022 | **0.063** | 0.020 | -0.011 | 0.024 | -0.050 | 0.024 | 0 |  |
| 10. Ka'a'awa | 0.157 | 0.168 | **0.147** | 0.217 | **0.221** | **0.235** | 0.142 | **0.223** | 0.164 | 0 |

**Figure 3.**
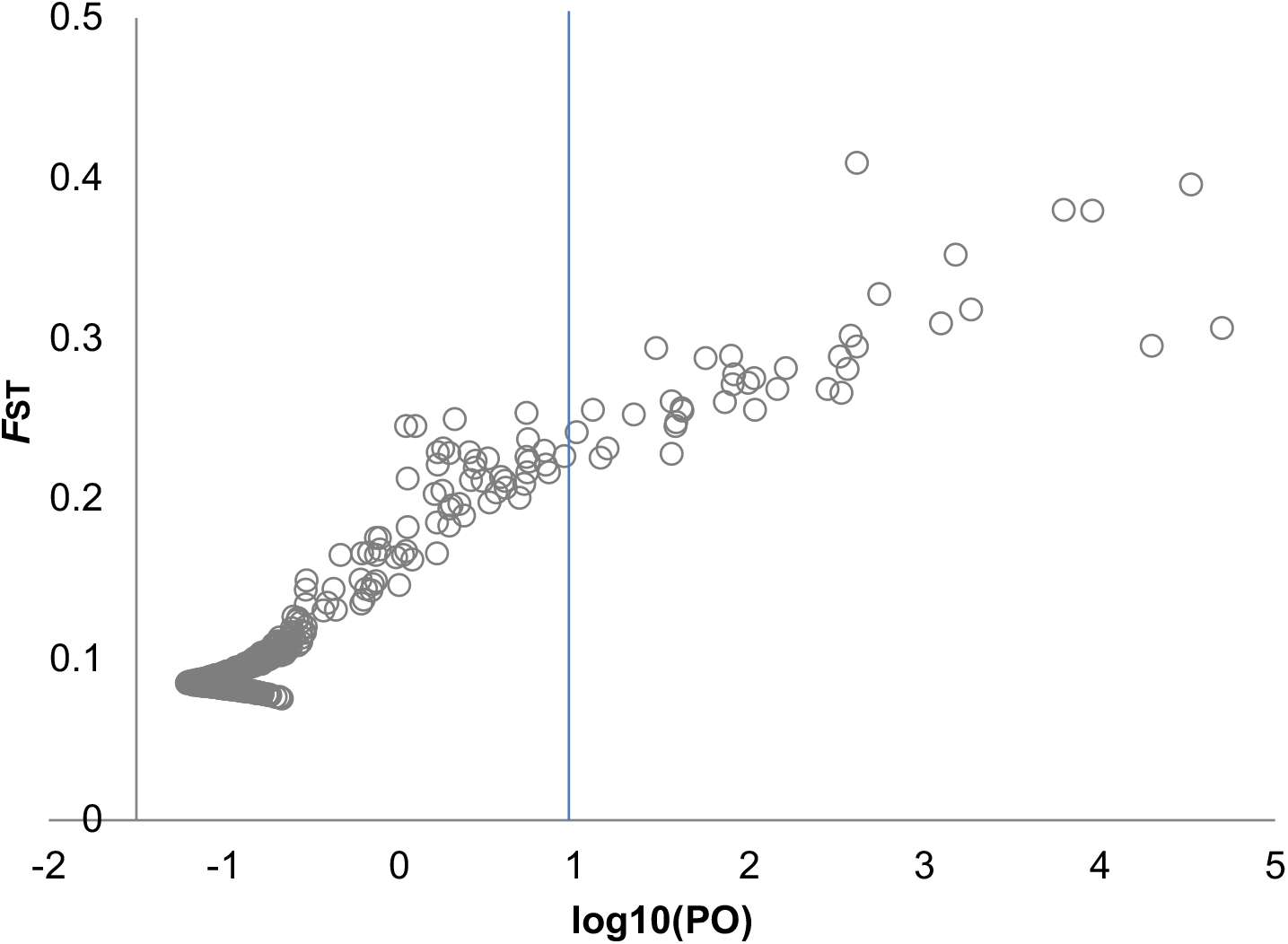
Outlier loci among sampled populations of *Coralliophila violacea as* identified by BayeScan. Points to the right of the blue line are outliers.

However, regional variation only accounted for 10% of the overall variation, and the majority (90%) was within populations (Table 4).

**Table 4.**
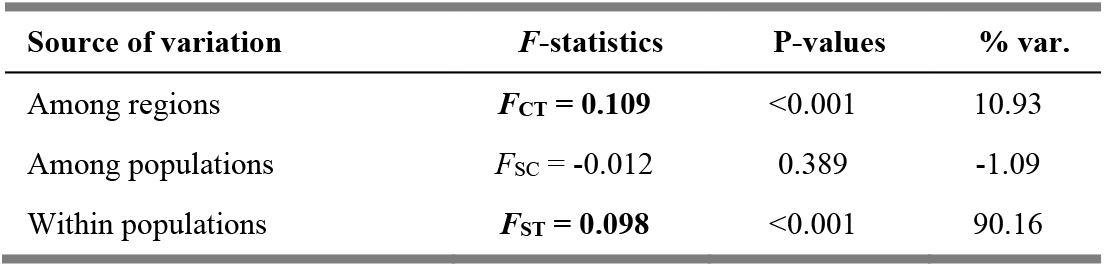
Hierarchical AMOVA of *Coralliophila violacea* populations sampled across all 3,188 loci with locations grouped by region as indicated in Table 1. Significant *F*-statistic values at *p* >0.001 are in boldface.

### Outlier tests

Using a false discovery rate (FDR) of 10%, BayeScan revealed 72 outlier loci putatively under directional selection among all locations (FDR = 0.10, 72; FDR = 0.05, 57), and 34 outlier loci among regions (Indian Ocean, Vietnam, Coral Triangle, Hawai i; FDR = 0.10, 34; FDR = 0.05, 25) (Fig. 4). Considering only these 72 outlier loci, STRUCTURE indicated that *K* = 2 was the best-supported number of genetic partitions (Fig. 2). In contrast to results from all loci or just neutral loci above, the two genetic partitions identified using only outlier loci do not correspond to individual sites. Instead, results recover two groups: 1) Hòn Mun, Vietnam and Ka’a’awa, Hawai i, two locations separated by ∼9,800 km of open ocean, and 2) all other sites (Fig. 2).

**Figure 4.**
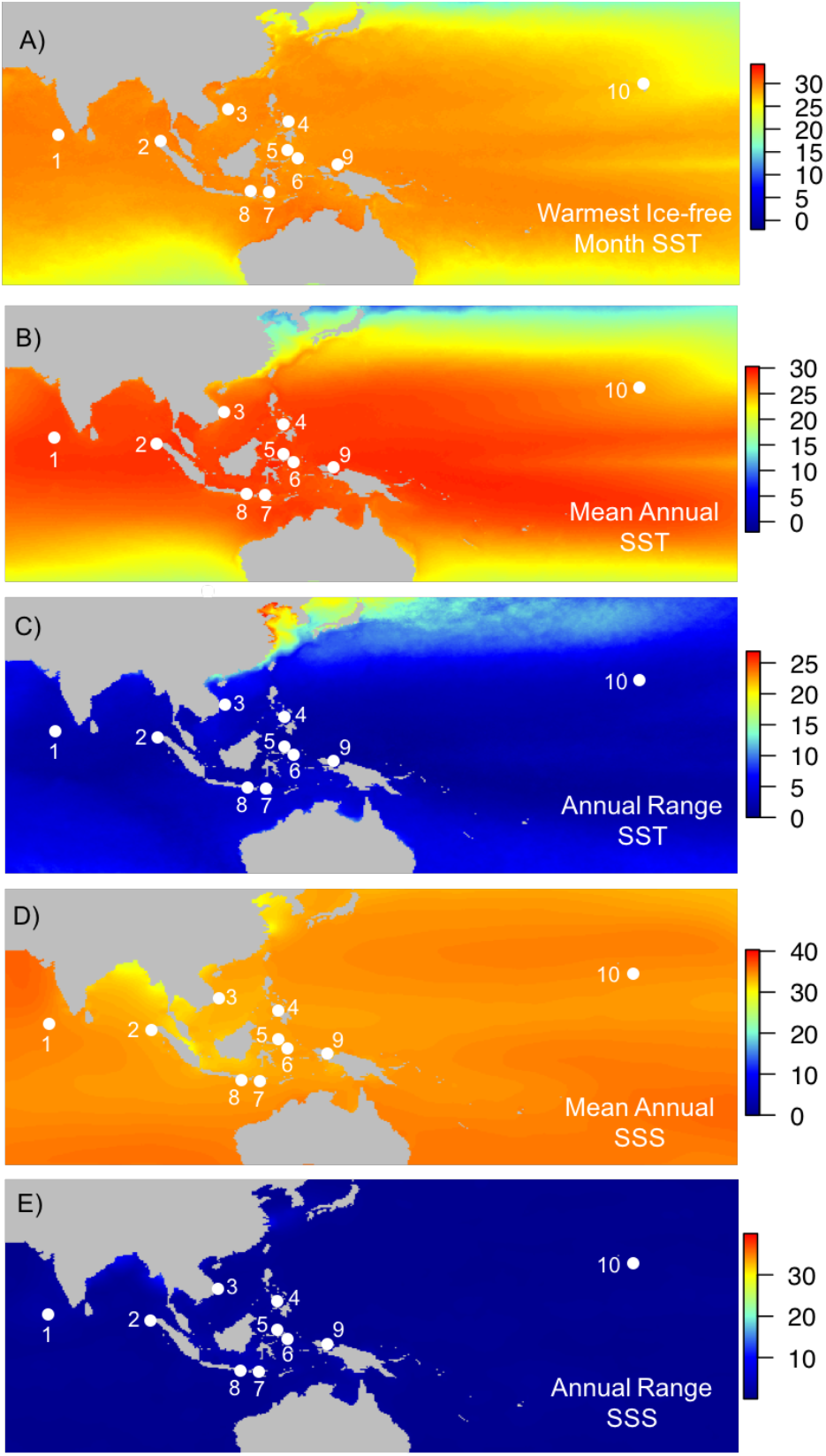
Ocean climate variables **a)** temperature of the warmest ice-free month, **b)** annual mean, and **c)** range of sea surface temperature (SST) in °C, **d)** annual mean, and **e)** range of sea surface salinity (SSS) in psu. Data source MARSPEC (Sbrocco & Barber, 2013).

### Ocean climate variables and associations with allele frequency differences

Results from MARSPEC show significant variation in the five environmental variables across the sampled region. The temperature of the warmest ice-free month varied by more than 3 °C across the sampled sites, with the highest temperature in the Indian Ocean (∼29.4 °C) and lowest in Hawai i (26.4 °C) and Vietnam (28.2 °C) (Table 1, Fig. 4a). The mean annual sea surface temperature (SST) also varied substantially across sampling locations by ∼4 °C (Table 1, Fig. 4a). However, in this case, Manokwari (28.90 °C) and Pulau Weh (28.59 °C) at the edges of Indonesia were the warmest, while again Hawai i (24.99 °C) and Vietnam (26.45 °C) were the coldest locations (Table 1, Fig. 4a). SST at some sites was very stable, with only small annual ranges: Manokwari (0.78 °C) and Bunaken (1.05 °C). At other locations, SST varied more widely (e.g., Dumaguete (3.16 °C) and Vietnam (4.22 °C); Table 1, Fig. 4b).

Sea surface salinity (SSS) also showed variation across sampling locations (Table 1, Fig. 2.5c). Sites in the two oceanic archipelagos, Vavvaru, Maldives (34.93 psu), and Ka’a’awa, Hawai i (34.92 psu), had the highest values. In contrast, salinity at sites in the Coral Triangle was consistently lower (33.89 – 34.09 psu) (Table 1, Fig. 4c). Locations also experienced annual variation in salinity with inputs from freshwater runoff and precipitation (Fig. 4d). Sites in the Indian Ocean, Vavvaru (SSS range = 1.64 psu), and Pulau Weh (SSS range = 1.43 psu) had the most variable salinity while Ka’a’awa (SSS range = 0.21 psu) in Hawai i was the most stable (Table 1, Fig. 4d).

We identified a total of 88 SNPs using Bayenv2, for which allele frequency differences correlated with one or more of the five ocean climate variables (Fig. 5). The greatest proportion of these 88 SNPs were associated with annual temperature range (*N* = 38), followed by mean annual temperature (*N* = 31), temperature of the warmest ice-free month (*N* = 21), annual range of salinity (*N* = 12) and mean annual salinity (*N* = 10) (Fig. 5). SNPs associated with temperature had the highest Bayes Factors (Fig. 5). Of these, 22 SNPs were associated with more than one variable (*N* = 20 with two variables, *N* = 2 with three variables). The warmest month’s mean SST and temperature were the most highly correlated, with 15 SNPs shared between them, followed by mean and range of SST (*N* = 5 SNPs). There was no overlap between the 88 loci associated with environmental variables and the 72 outlier loci.

**Figure 5.**
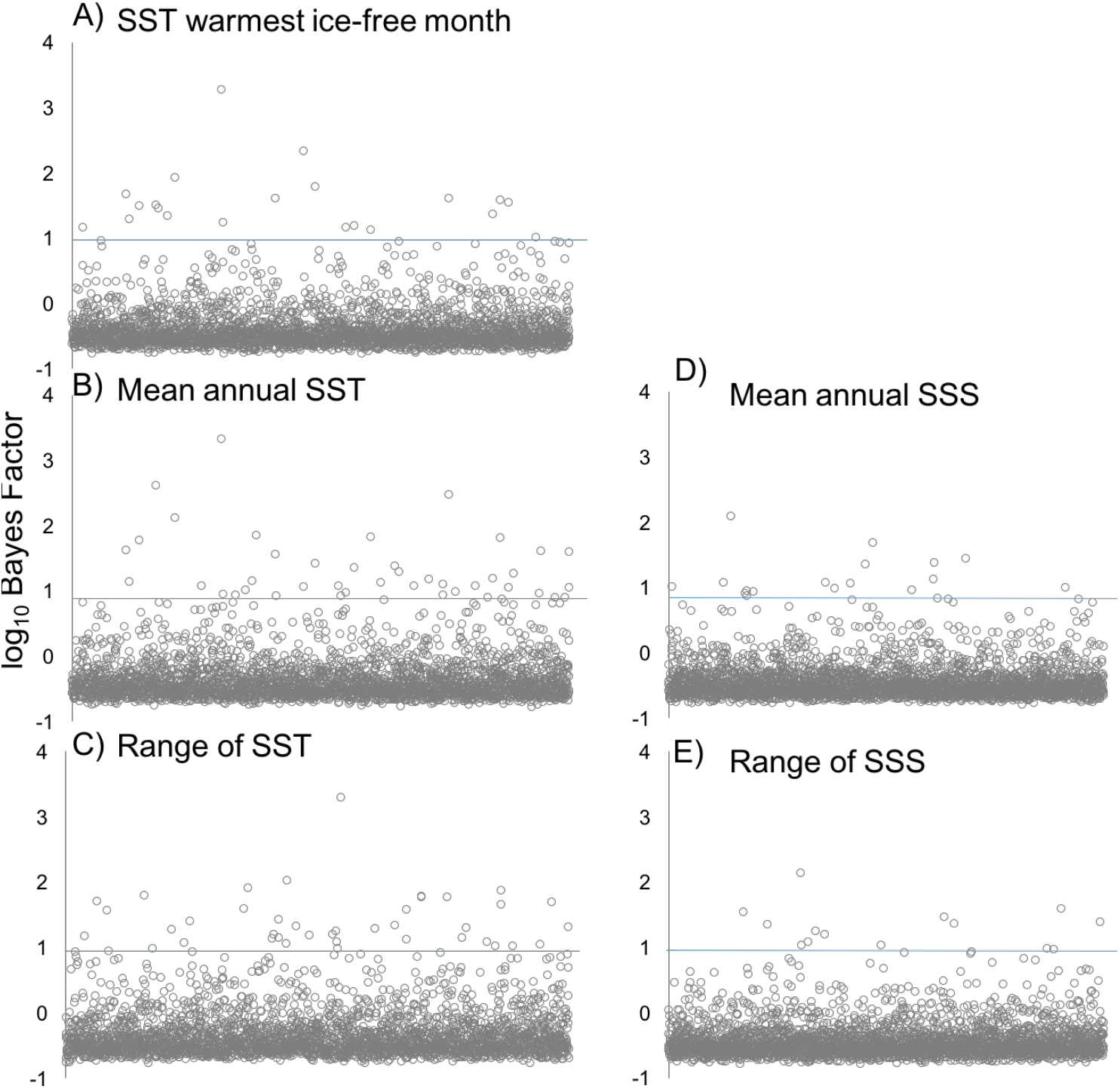
Bayenv2 association of allele frequencies for all polymorphic loci (3,186) with each ocean climate variable. SNPs are ordered by tagID. Blue line indicates significance level above which loci are considered to associate with climate variables (Jeffreys, 1998) strongly.

### Mapping and annotation of outliers and environmentally-associated loci

For loci significantly associated with environmental variables, the majority (49/87; 56%) did not map to any mollusc or lophotrochozoan sequences available in the NCBI database (Table 5). However, we were able to align the other 43% (38/88) of candidate loci to DNA sequences from a variety of molluscs, including an octopus (*Octopus vulgaris*), oysters (*Crassostrea gigas, C. virginica*), scallops (*Pecten maximus, Mizuhopecten yessoensis, Azumapecten farrei*), a clam (*Scapharca broughtonii*), and four marine snails (*Conus episcopatus, C. ximenes, Thais clavigera, Nucella lapillus*) and two freshwater snails (*Pomacea canaliculata, Biomphalaria glabrata*). Of these loci, 19 loci were mapped to protein-coding regions. Six of these loci coded for uncharacterized proteins, two microsatellites, and the other 11 were characterized with annotated functions (Table 5).

**Table 5.**
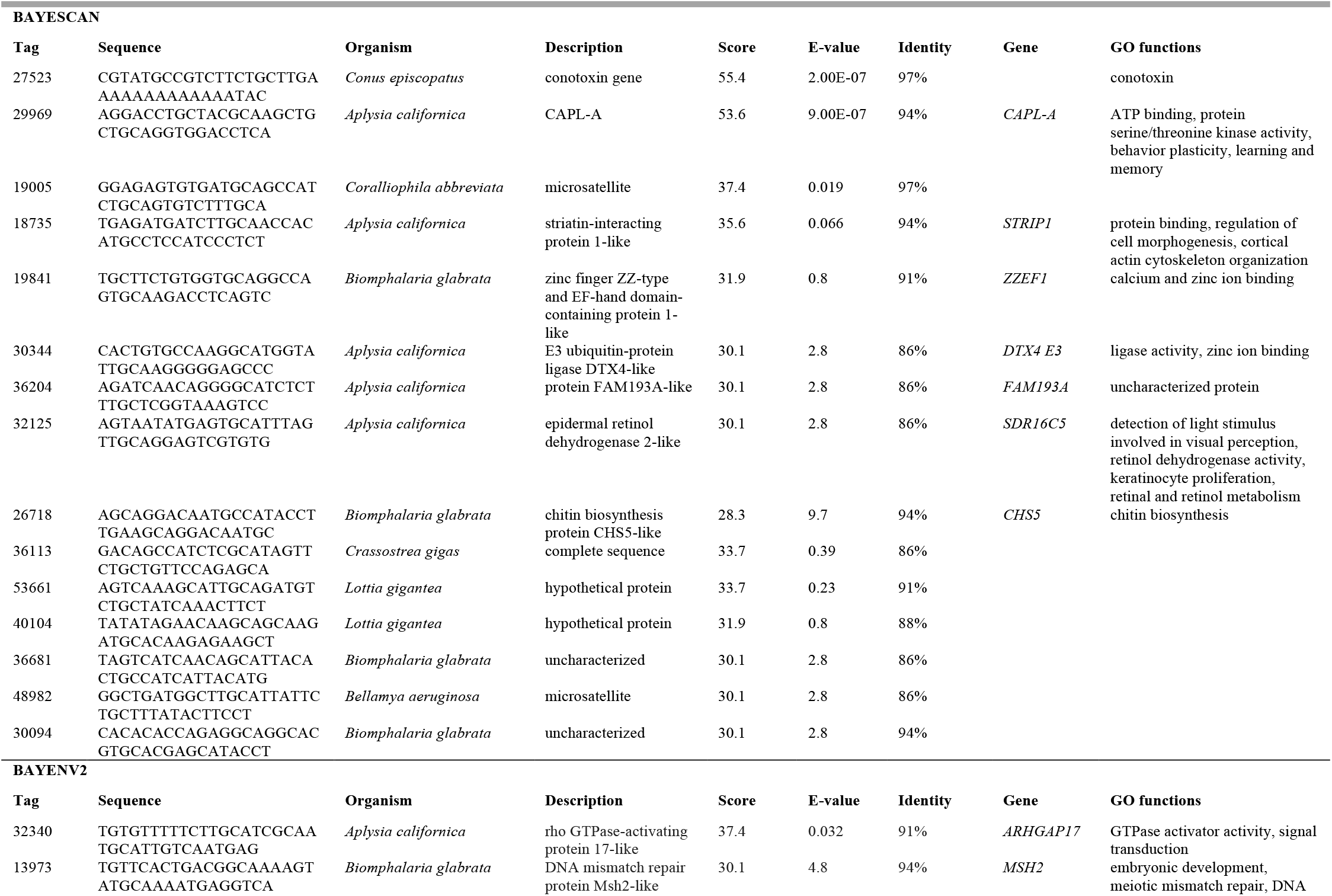
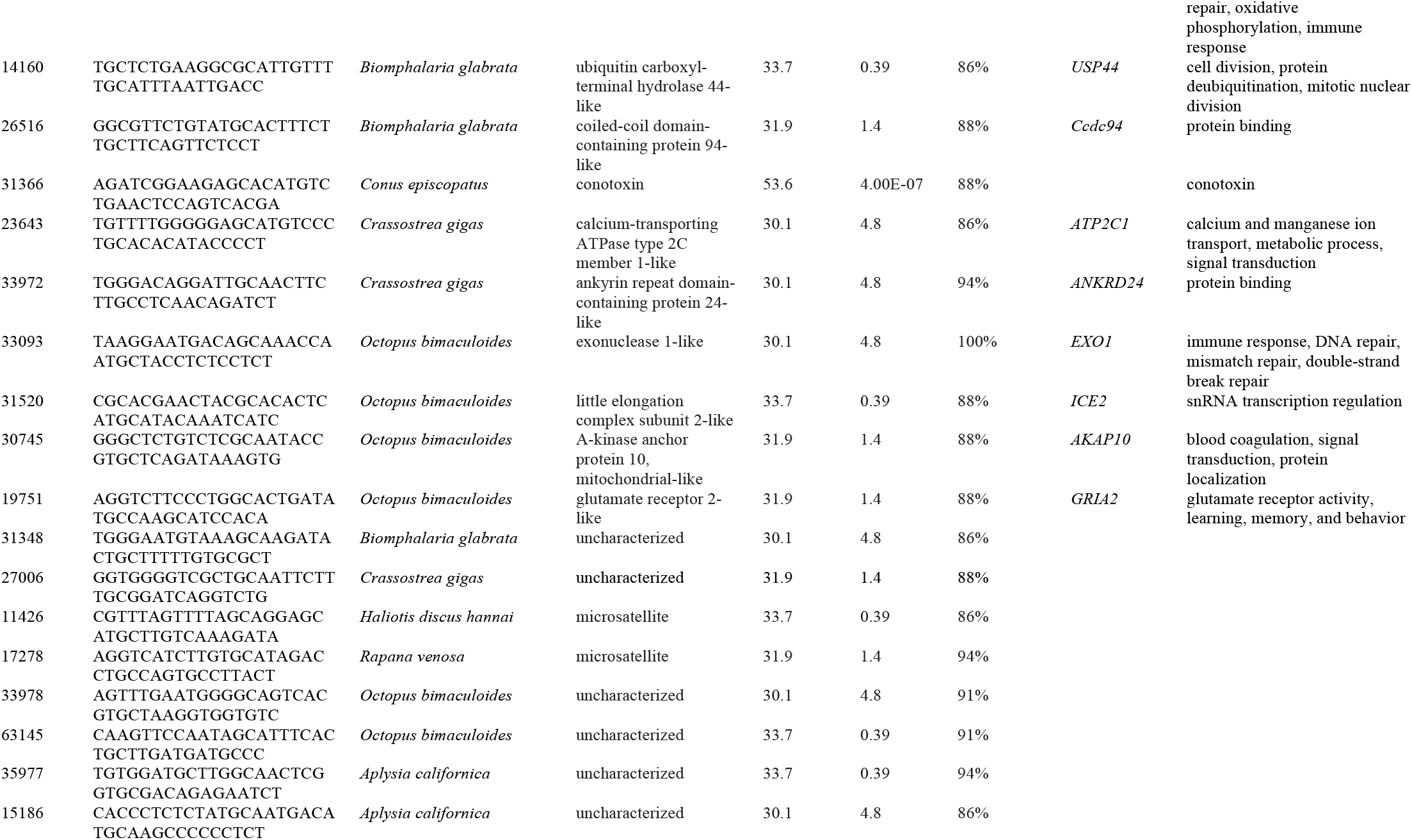
Annotations for BayeScan outlier loci among sites and Bayenv2 environmentally-associated loci mapped to DNA sequences in the NCBI database.

For outlier loci identified by BayeScan, nearly half (47%; 34/72) mapped to molluscs in the NCBI database, including an octopus (*Octopus pallidus*), oysters (*Crassostrea gigas, C. virginica*), a scallop (*Pecten maximus*), a sea hare (*Aplysia californica*), a limpet (*Lottia gigantea*), a freshwater snail (*Pomacea canaliculata*) and marine snail (*Conus episcopatus*) (Table 5). Of these loci, ten had associated predicted protein functions. The top two outlier loci, in terms of percent identity of sequences and the lowest E-value, aligned with a conotoxin from a cone snail (*Conus episcopatus*) and a gene (*CAPLA*) coding for a protein involved in behavioral plasticity in the California seahare (*Aplysia californica*). Interestingly, despite no overlap between the 72 outlier loci identified by BayeScan and the 88 environmentally-associated loci identified by Bayenv2, both sets contained loci mapped to genes coding for conotoxins (Table 5).

## DISCUSSION

Our results reveal a complex mixture of population genetic patterns in *Coralliophila violacea* that appear to be shaped by both geography and environment. Genome-wide data from >3,000 SNPs identified four genetic partitions concordant with traditionally recognized biogeographic regions (Indian Ocean, South China Sea, Coral Triangle, and Hawai i; (Spalding et al., 2007). These findings align with a previous mitochondrial DNA study (Simmonds et al., 2018) and other marine phylogeographic studies from this region (Barber et al., 2011; Carpenter et al., 2010; Crandall et al., 2019). Furthermore, repeated patterns across multiple taxa suggest a common origin (Avise 2000), the most likely broad-scale physical isolation of populations— allopatry.

However, an examination of outlier loci and the correlation between allelic variation and environmental variables also highlights the importance of natural selection in shaping population genetic patterns. More than 2% of loci were outliers, similar to Gaither et al. (2015), showing that divergence is greater than expected under neutrality (Foll & Gaggiotti, 2008). Similarly, nearly 3% of loci had strong correlations with environmental variables. Both Hawai i and Vietnam, peripheral populations with similar climate profiles, were differentiated at neutral and outlier loci. In contrast, populations more centrally located in the range of *C. violacea* were only differentiated at neutral loci but not outliers. These results suggest that environmental differences, particularly sea surface temperature, may reinforce the neutral geographic structure, leading to peripheral populations’ differentiation. Combined with the ecological differentiation of *C. violacea* on different coral hosts (Simmonds et al., 2020), these results indicate that diversification in marine environments is more complex and varied than currently understood and includes both neutral and selective processes.

### Potential role of natural selection in shaping genetic patterns

While neutral loci showed a clear signal of physical limits to gene flow (below), there were also significant associations between frequencies of multiple loci (3% of total) and environmental variables. The strongest genetic associations linked these loci to the SST range and SST mean. Elevated SST can affect gastropods’ growth and survival rates, including marine snails (Sanford & Kelly, 2011), so the link between environmentally-associated loci and temperature is not unexpected. However, results also showed an interesting genetic association pattern driven by the temperature at the coldest sites in the peripheral populations Hawai i and Vietnam. Thus, temperature differences between the Coral Triangle and peripheral populations could enable selection to drive population divergence. For example, Reid et al. (2006) found that environmental and ecological factors (i.e., continental vs. oceanic habitat and primary productivity) shaped intertidal snails’ genetics in the Indo-Pacific. Given that peripheral populations like Hawai i have environmental conditions considered marginal for *Porites*’ growth and survival (Polato et al., 2010), such selection could act directly on *C. violacea* or indirectly by selection pressure on their coral hosts. Interestingly, while all four regions sampled show unique environmental signatures when examining the loci under the environmental influence, STRUCTURE only differentiates between two clusters: Vietnam and Hawai i to the exclusion of all others.

If environmental variation drives natural selection, it is reasonable to expect some overlap between loci with significant associations with environmental variables and the outlier loci that are putatively under selection. However, there was no overlap between these two groups. This seemingly incongruous result may be a function of only examining five environmental variables or potentially due to uneven sample sizes. We focused on temperature and salinity because of previous studies demonstrating their selection pressure on marine populations (e.g., Berg et al., 2015; DeFaveri et al., 2013; Limborg et al., 2012; Teske et al., 2019). However, other variables can have an impact. For example, Gaither et al. (2015) found that high turbidity and low light conditions explained divergent selection at a visual pigment gene locus in a coral reef fish.

As expected for a non-model organism, few candidate loci aligned to sequences in the NCBI database, highlighting the lack of current genomic resources for gastropods. However, two groups of genes are of interest for future research. First, conotoxin genes showed up as both outlier and environmentally-associated loci. While neurotoxins have not been studied in *Coralliophila*, venoms have been identified in other Neogastropoda (Fänge, 1984) and could be important to deactivate nematocysts to facilitate feeding and living on corals. Second, another gene of interest, *CHS5*, is predicted to be involved in chitin biosynthesis, a substance found in mollusc shells and radulae. Chitin is also used in the epidermis and stomach lining of nudibranchs that feed on Cnidarians as protection from nematocysts (Martin et al., 2007), and *Coralliophila* could similarly deploy chitin.

### Populations at the periphery

Hawaiian populations of *C. violacea* were the most geographically isolated, ∼7000 km removed from the Coral Triangle, and showed the strongest genetic differentiation. Studies reporting genetic differentiation of Hawaiian populations in Indo-Pacific taxa are common and frequently invoke geographic isolation (Ahti et al., 2016; J. DiBattista et al., 2012; Fernandez-Silva et al., 2015; Leray et al., 2010; Waldrop et al., 2016), and population bottlenecks (Szab’o et al., 2014) as primary drivers. Not only are Hawaiian populations of *C. violacea* differentiated, so too is their host, *Porites lobata* (Baums et al., 2012; Polato et al., 2010), suggesting a common diversification process. Whether by pure allopatry or isolation by distance, there are physical limits to gene flow in *C. violacea* across this vast span of the ocean. In contrast, Vietnam populations are much less geographically isolated from the core of the geographic range of *C. violacea*. Our sampling location was only 10s to 100s of km from other reef systems in the South China Sea with high connectivity to the Coral Triangle (Treml et al., 2012). Such extreme geographic isolation cannot explain the differentiation of Vietnam.

Allopatric processes may contribute to the strong differentiation of *C. violacea* populations from Vietnam. The South China Sea was partially enclosed during Plio-Pleistocene low sea levels (Ludt & Rocha, 2015; Voris, 2000), resulting in isolation and genetic differentiation of marine taxa (Barber et al., 2000). Ocean circulation patterns also limit water movement from the Coral Triangle into the South China Sea (Kool et al., 2011), creating a barrier dividing populations north and south across the middle of the South China Sea (Treml et al., 2015). Ocean circulation patterns have been invoked to explain *C. violacea* populations’ genetic structure from Taiwan and Taiping, a small island in the South China Sea (Lin & Liu, 2008). These allopatric processes may fully explain differentiation at neutral loci. However, they can’t explain the grouping of Hawai i and Vietnam populations when examining loci exhibiting signals of selection. The similarity of these populations at non-neutral loci, combined with their environmental similarities, strongly argues that environmental variation drives or reinforces differentiation of these peripheral populations.

Interestingly, while peripheral populations in the Indian Ocean (Vavvaru and Pulau Weh) showed a distinct Indian Ocean signature, these populations also had a substantial admixture generating from the Coral Triangle and the South China Sea. This is not surprising given the volume of sea surface transport from the Pacific into the Indian Ocean through this region (Gordon et al., 2012), a pattern that is observed in a wide variety of taxa (e.g., Drew & Barber, 2009; Williams & Benzie, 1998). However, this region’s environmental conditions are also different from other areas of the sampled range, suggesting that selection might act as it does in other parts of the range of *C. violacea*. Thus, gene flow from the Coral Triangle and the South China Sea or other un-sampled areas may limit the signal of environmental selection in the Maldives, despite the relative isolation of these reef ecosystems.

### Populations at the center

In contrast to peripheral populations, populations within the Coral Triangle at the core of *C. violacea’s* geographic range showed no evidence of divergence. Populations in the Philippines (Dumaguete), eastern (Manokwari), and central Indonesia (Bunaken, Lembeh, Pulau Mengyatan, Nusa Penida) all had high connectivity, echoing findings for other molluscs in the region (S. H. Cheng et al., 2014; DeBoer et al., 2014; Kirkendale & Meyer, 2004; Nuryanto & Kochzius, 2009) and as predicted by coupled biophysical models of larval dispersal (Kool et al., 2011; Treml et al., 2015). The only isolation observed by Coral Triangle populations was from populations on the periphery in the South China Sea, the Indian Ocean, and the Central Pacific.

High connectivity, however, is not expected among all populations in the Coral Triangle. Specifically, studies of multiple taxa including giant clams (DeBoer et al., 2008, 2014; Kochzius & Nuryanto, 2008); mantis shrimp (Barber et al., 2006, 2011); echinoderms (Crandall et al., 2008b); and fish (Timm & Kochzius, 2008; Ackiss et al., 2013; Jackson et al., 2014) all show strong isolation of populations spanning the Maluku Sea. This pattern has typically been attributed to the effects of the Halmahera Eddy (Barber et al., 2006) and is predicted by coupled biophysical models (Kool et al., 2011; Treml et al., 2015). However, Treml et al. (2015) found that the Halmahera Eddy filtered only 8.3 – 10.6% of simulated taxa. The strength of this filter was determined by life-history traits such as reproductive output, the timing of spawning, and length of larval dispersal phase. For example, *C. violacea* has high fecundity (Lin & Liu, 1995), spawn larvae most of the year (Soong & Chen, 1991), and likely have a long larval duration (Taylor, 1975). These traits should facilitate crossing seasonally variable dispersal barriers. In contrast, giant clams (*Tridacna*) have relatively short-lived planktonic larvae (9–10 days; Crawford et al., 1986), constraining their dispersal potential, making them more inclined to diverge across filters to gene flow. As such, life history difference likely explains why connectivity in the Coral Triangle isn’t limited in *C. violacea*, as it is in other molluscs.

While allopatric divergence is an important process shaping marine taxa’s evolutionary history, physical limits to dispersal and gene flow do not operate in isolation. Our results show that local adaptation to different environments may reinforce neutral divergence, especially in peripheral populations. These results add to the growing recognition that various factors impact diversification in coral reef species (Bowen et al., 2013; Fernandez-Silva et al., 2015; Floeter et al., 2007; Hoeksema, 2007; Ludt & Rocha, 2015; Rocha & Bowen, 2008). Advances in high throughput genome-wide sequencing will facilitate the exploration of neutral and adaptive variation in concert. Combined with the improved availability of marine environmental databases, we are at the leading edge of developing a more expansive view of processes driving genetic divergence among populations, leading to clearer approaches to understanding evolution in marine habitats.

## ACKNOWLEDGMENTS

This work was supported by three National Science Foundation programs (OISE-0730256, OISE-1243541, and OCE-0349177) and a US Agency for International Development Grant (497-A-00-10-00008-00). The Lemelson Foundation Fellowship, Conchologists of America, Sigma Xi, and the UCLA Department of Ecology and Evolutionary Biology provided additional funding to S. Simmonds. Finally, we acknowledge support from the Indonesian government, including the Indonesian Ministry of Research and Technology (RISTEK), the Indonesian Institute of Sciences (LIPI), the Nature Conservation Agency (BKSDA), and the National Marine Park offices of Bunaken and Wakatobi. Sampling was covered under research permits obtained in Indonesia (RISTEK 2011, 198/SIP/FRP/SMNl/2012, 187/SIP/FRP/SM/VI/2013), Timor-Leste (Direccao Nacional de Pescase Aquicultura 0042/DNPA/IOP/VII/11), Vietnam, the Philippines (Department of Agriculture-Bureau of Fisheries and Aquatic Resources), the Maldives (Ministry of Fisheries and Aquaculture Permit No. (OTHR)30-D/INDIV/2013/116) and Hawai i (Department of Land and Natural Resources SAP 2013-11). We also thank the Indonesian Biodiversity Research Center at Udayana University, Institute for Environmental and Marine Sciences at Silliman University, and Nha Trang University for institutional support.

Thank you to B. Stockwell, M. Weber, H. Nuetzel, and D. Willette for collecting specimens.

## Author Contributions

SES and PHB secured funding, conceptualized the study, worked out the methodology, and led the original manuscript’s writing. SES, AFP, SHC, and PHB obtained permits and collected samples. SES, AFP, and SHC conducted lab work. SES performed data analyses and visualization. GM provided resources and project administration. All co-authors contributed to editing and revisions.

## REFERENCES

Ackiss, A. S., Bird, C. E., Akita, Y., Santos, M. D., Tachihara, K., & Carpenter, K. E. (2018). Genetic patterns in peripheral marine populations of the fusilier fish Caesio cuning within the Kuroshio Current. Ecology and Evolution, 8(23), 11875–11886. https://doi.org/10.1002/ece3.4644

Ackiss, A. S., Pardede, S., Crandall, E. D., Ablan-Lagman, M. C. A., Ambariyanto Romena, N., Barber, P. H., & Carpenter, K. E. (2013). Pronounced genetic structure in a highly mobile coral reef fish, Caesio cuning, in the Coral Triangle. Marine Ecology Progress Series, 480, 185–197. https://doi.org/10.3354/meps10199

Ahti, P. A., Coleman, R. R., DiBattista, J. D., Berumen, M. L., Rocha, L. A., & Bowen, B. W. (2016). Phylogeography of Indo-Pacific reef fishes: sister wrasses Coris gaimard and C. cuvieri in the Red Sea, Indian Ocean and Pacific Ocean. Journal of Biogeography, 43(6), 1103–1115. https://onlinelibrary.wiley.com/doi/abs/10.1111/jbi.12712

Barber, P.H., & Meyer, C.P. (2014) Pluralism explains diversity in the Coral Triangle. In: Mora C (ed) Ecology of Fishes on Coral Reefs: pp. 258–263; Cambridge University Press.

Barber, P. H., Cheng, S. H., Erdmann, M. V., & Tengardjaja, K. (2011). Evolution and conservation of marine biodiversity in the Coral Triangle: insights from stomatopod Crustacea. Crustacean Issues, 19, 129–156. https://content.taylorfrancis.com/books/e/download?dac=C2010-0-27653-7&isbn=9781439840740&doi=10.1201/b11113-10&format=pdf

Barber, P. H., Erdmann, M. V., & Palumbi, S. R. (2006). Comparative phylogeography of three codistributed stomatopods: origins and timing of regional lineage diversification in the Coral Triangle. Evolution, 60(9), 1825–1839. https://www.ncbi.nlm.nih.gov/pubmed/17089967

Barber, P. H., Palumbi, S. R., Erdmann, M. V., & Moosa, M. K. (2000). Biogeography. A marine Wallace’s line? Nature, 406(6797), 692–693. https://doi.org/10.1038/35021135

Baums, I. B., Boulay, J. N., Polato, N. R., & Hellberg, M. E. (2012). No gene flow across the Eastern Pacific Barrier in the reef-building coral Porites lobata. Molecular Ecology, 21(22), 5418–5433. https://doi.org/10.1111/j.1365-294X.2012.05733.x

Berg, P. R., Jentoft, S., Star, B., Ring, K. H., Knutsen, H., Lien, S., Jakobsen, K. S., & André, C. (2015). Adaptation to low salinity promotes genomic divergence in Atlantic Cod (Gadus morhua L.). Genome Biology and Evolution, 7(6), 1644–1663. https://doi.org/10.1093/gbe/evv093

Bowen, B. W., Rocha, L. A., Toonen, R. J., Karl, S. A., & ToBo Laboratory. (2013). The origins of tropical marine biodiversity. Trends in Ecology & Evolution, 28(6), 359–366. https://doi.org/10.1016/j.tree.2013.01.018

Bowen, B. W., Shanker, K., Yasuda, N., Celia, M., Malay, M. C. (machel) D., von der Heyden, S., Paulay, G., Rocha, L. A., Selkoe, K. A., Barber, P. H., Williams, S. T., Lessios, H. A., Crandall, E. D., Bernardi, G., Meyer, C. P., Carpenter, K. E., & Toonen, R. J. (2014). Phylogeography unplugged: comparative surveys in the genomic era. Bulletin of Marine Science, 90(1), 13–46. https://doi.org/10.5343/bms.2013.1007

Briggs, J. C. (1992). The marine East Indies: Centre of origin? Global Ecology and Biogeography Letters, 2(5), 149. https://doi.org/10.2307/2997803

Briggs, J. C. (2006). Proximate sources of marine biodiversity. Journal of Biogeography, 33(1), 1–10. https://doi.org/10.1111/j.1365-2699.2005.01374.x

Burgess, S. C., Treml, E. A., & Marshall, D. J. (2012). How do dispersal costs and habitat selection influence realized population connectivity? Ecology, 93(6), 1378–1387. https://www.ncbi.nlm.nih.gov/pubmed/22834378

Carpenter, K. E., Barber, P. H., Crandall, E. D., Ablan-Lagman, M. C. A., Ambariyanto Mahardika, G. N., Manjaji-Matsumoto, B. M., Juinio-Meñez, M. A., Santos, M. D., Starger, C. J., & Toha, A. H. A. (2010). Comparative phylogeography of the Coral Triangle and Implications for Marine Management. Journal of Marine Biology, 2011. https://doi.org/10.1155/2011/396982

Cheng, S. H. (2015). Evolution and Population Genomics of Loliginid Squids [UCLA]. https://escholarship.org/uc/item/0zw3h4ps

Cheng, S. H., Anderson, F. E., Bergman, A., Mahardika, G. N., Muchlisin, Z. A., Dang, B. T., Calumpong, H. P., Mohamed, K. S., Sasikumar, G., Venkatesan, V., & Barber, P. H. (2014). Molecular evidence for co-occurring cryptic lineages within the Sepioteuthis cf. lessoniana species complex in the Indian and Indo-West Pacific Oceans. Hydrobiologia, 725(1), 165–188. https://doi.org/10.1007/s10750-013-1778-0

Coop, G., Witonsky, D., Di Rienzo, A., & Pritchard, J. K. (2010). Using environmental correlations to identify loci underlying local adaptation. Genetics, 185(4), 1411–1423. https://doi.org/10.1534/genetics.110.114819

Craig, M. T., Eble, J. A., Bowen, B. W., & Robertson, D. R. (2007). High genetic connectivity across the Indian and Pacific Oceans in the reef fish Myripristis berndti (Holocentridae). Marine Ecology Progress Series, 334, 245–254. https://doi.org/10.3354/meps334245

Crandall, E. D., Frey, M. A., Grosberg, R. K., & Barber, P. H. (2008a). Contrasting demographic history and phylogeographical patterns in two Indo-Pacific gastropods. Molecular Ecology, 17(2), 611–626. https://doi.org/10.1111/j.1365-294X.2007.03600.x

Crandall, E. D., Jones, M. E., Muñoz, M. M., Akinronbi, B., Erdmann, M. V., & Barber, P. H. (2008b). Comparative phylogeography of two seastars and their ectosymbionts within the Coral Triangle. Molecular Ecology, 17(24), 5276–5290. https://doi.org/10.1111/j.1365-294X.2008.03995.x

Crandall, E. D., Sbrocco, E. J., DeBoer, T. S., Barber, P. H., & Carpenter, K. E. (2012) Expansion dating: Calibrating molecular clocks in marine species from expansions onto the Sunda Shelf following the last glacial maximum, Molecular Biology and Evolution, 29(2) 707–719, https://doi.org/10.1093/molbev/msr227

Crandall, E. D., Riginos, C., Bird, C., Liggins, L., Treml, E., Beger, M., Barber, P. H., Connolly, S. R., Cowman, P. F., DiBattista, J. D., Eble, J. A., Magnuson, S. F., Horne, J. B., Kochzius, M., Lessios, H. A., Yin Vanson Liu, S., Ludt, W. B., Madduppa, H., Pandolfi, J. M., … Gaither, M. R. (2019). The molecular biogeography of the Indo-Pacific: testing hypotheses with multispecies genetic patterns. Global Ecology and Biogeography: A Journal of Macroecology. https://onlinelibrary.wiley.com/doi/abs/10.1111/geb.12905

Crawford, C. M., Nash, W. J., & Lucas, J. S. (1986). Spawning induction, and larval and juvenile rearing of the giant clam, Tridacna gigas. Aquaculture, 58(3-4), 281–295. https://doi.org/10.1016/0044-8486(86)90094-3

DeBoer, T. S., Naguit, M. R. A., Erdmann, M. V., Ablan-Lagman, M. C. A., Ambariyanto Carpenter, K. E., Toha, A. H. A., & Barber, P. H. (2014). Concordance between phylogeographic and biogeographic boundaries in the Coral Triangle: Conservation implications based on comparative analyses of multiple giant clam species. Bulletin of Marine Science, 90(1), 277–300. https://doi.org/10.5343/bms.2013.1003

DeBoer, T. S., Subia, M. D., Ambariyanto Erdmann, M. V., Kovitvongsa, K., & Barber, P. H. (2008). Phylogeography and limited genetic connectivity in the endangered boring giant clam across the Coral Triangle. Conservation Biology, 22(5), 1255–1266. https://doi.org/10.1111/j.1523-1739.2008.00983.x

DeFaveri, J., Jonsson, P. R., & Merilä, J. (2013). Heterogeneous genomic differentiation in marine threespine sticklebacks: adaptation along an environmental gradient. Evolution, 67(9), 2530–2546. https://doi.org/10.1111/evo.12097

DiBattista, J. D., Roberts, M. B., Bouwmeester, J., Bowen, B. W., Coker, D. J., Lozano-Cortés, D. F., Howard Choat, J., Gaither, M. R., Hobbs, J.-P. A., Khalil, M. T., & Others. (2016). A review of contemporary patterns of endemism for shallow water reef fauna in the Red Sea. Journal of Biogeography, 43(3), 423–439. https://onlinelibrary.wiley.com/doi/abs/10.1111/jbi.12649

DiBattista, J., Waldrop, E., Bowen, B. W., Schultz, J. K., Gaither, M. R., Pyle, R. L., & Rocha, L. A. (2012). Twisted sister species of pygmy angelfishes: discordance between taxonomy, coloration, and phylogenetics. Coral Reefs, 31(3), 839–851. https://doi.org/10.1007/s00338-012-0907-y

Dohna, T. A., Timm, J., Hamid, L., & Kochzius, M. (2015). Limited connectivity and a phylogeographic break characterize populations of the pink anemonefish, Amphiprion perideraion, in the Indo-Malay Archipelago: inferences from a mitochondrial and microsatellite loci. Ecology and Evolution, 5(8), 1717–1733. https://doi.org/10.1002/ece3.1455

Drew, J., & Barber, P. H. (2009). Sequential cladogenesis of the reef fish Pomacentrus moluccensis (Pomacentridae) supports the peripheral origin of marine biodiversity in the Indo-Australian archipelago. Molecular Phylogenetics and Evolution, 53(1), 335–339. https://doi.org/10.1016/j.ympev.2009.04.014

Evanno, G., Regnaut, S., & Goudet, J. (2005). Detecting the number of clusters of individuals using the software STRUCTURE: a simulation study. Molecular Ecology, 14(8), 2611–2620. https://doi.org/10.1111/j.1365-294X.2005.02553.x

Excoffier, L., & Lischer, H. E. L. (2010). Arlequin suite ver 3.5: a new series of programs to perform population genetics analyses under Linux and Windows. Molecular Ecology Resources, 10(3), 564–567. https://doi.org/10.1111/j.1755-0998.2010.02847.x

Fänge, R. (1984). Venoms and venom glands of marine molluscs. In Bolis L., Zadunaisky J., Gilles R. (Ed.), Toxins, Drugs, and Pollutants in Marine Animals (pp. 47–62). Springer Berlin Heidelberg. https://doi.org/10.1007/978-3-642-69903-0_5

Fernandez-Silva, I., Randall, J. E., Coleman, R. R., DiBattista, J. D., Rocha, L. A., Reimer, J. D., Meyer, C. G., & Bowen, B. W. (2015). Yellow tails in the Red Sea: phylogeography of the Indo-Pacific goatfish Mulloidichthys flavolineatus reveals isolation in peripheral provinces and cryptic evolutionary lineages. Journal of Biogeography, 42(12), 2402–2413. https://onlinelibrary.wiley.com/doi/abs/10.1111/jbi.12598

Floeter, S. R., Rocha, L. A., Robertson, D. R., Joyeux, J. C., Smith-Vaniz, W. F., Wirtz, P., Edwards, A. J., Barreiros, J. P., Ferreira, C. E. L., Gasparini, J. L., Brito, A., Falcón, J. M., Bowen, B. W., & Bernardi, G. (2007). Atlantic reef fish biogeography and evolution. Journal of Biogeography, 35(1), 22–47. https://doi.org/10.1111/j.1365-2699.2007.01790.x

Foll, M., & Gaggiotti, O. (2008). A genome-scan method to identify selected loci appropriate for both dominant and codominant markers: a Bayesian perspective. Genetics, 180(2), 977–993. https://doi.org/10.1534/genetics.108.092221

Forsman, Z. H., Barshis, D. J., Hunter, C. L., & Toonen, R. J. (2009). Shape-shifting corals: molecular markers show morphology is evolutionarily plastic in Porites. BMC Evolutionary Biology, 9, 45. https://doi.org/10.1186/1471-2148-9-45

Forsman, Z., Wellington, G. M., Fox, G. E., & Toonen, R. J. (2015). Clues to unraveling the coral species problem: distinguishing species from geographic variation in Porites across the Pacific with molecular markers and microskeletal traits. PeerJ, 3, e751. https://doi.org/10.7717/peerj.751

Gaither, M. R., Bernal, M. A., Coleman, R. R., Bowen, B. W., Jones, S. A., Simison, W. B., & Rocha, L. A. (2015). Genomic signatures of geographic isolation and natural selection in coral reef fishes. Molecular Ecology, 24(7), 1543–1557. https://doi.org/10.1111/mec.13129

Gaither, M. R., Bowen, B. W., Bordenave, T.-R., Rocha, L. A., Newman, S. J., Gomez, J. A., van Herwerden, L., & Craig, M. T. (2011). Phylogeography of the reef fish Cephalopholis argus (Epinephelidae) indicates Pleistocene isolation across the Indo-Pacific barrier with contemporary overlap in the Coral Triangle. BMC Evolutionary Biology, 11(1), 189. https://doi.org/10.1186/1471-2148-11-189

Gaither, M. R., Toonen, R. J., Robertson, D. R., Planes, S., & Bowen, B. W. (2010). Genetic evaluation of marine biogeographical barriers: perspectives from two widespread Indo-Pacific snappers (Lutjanus kasmira and Lutjanus fulvus). Journal of Biogeography, 37(1), 133–147. https://onlinelibrary.wiley.com/doi/abs/10.1111/j.1365-2699.2009.02188.x

Gavrilets, S. (2003). Perspective: models of speciation: what have we learned in 40 years? Evolution; International Journal of Organic Evolution, 57(10), 2197–2215. https://onlinelibrary.wiley.com/doi/abs/10.1111/j.0014-3820.2003.tb00233.x

Gordon, A. L., Huber, B. A., & Metzger, E. J. (2012). South China Sea throughflow impact on the Indonesian throughflow. Geophysical Research Letters, 39(11).

Günther, T., & Coop, G. (2013). Robust identification of local adaptation from allele frequencies. Genetics, 195(1), 205–220. https://doi.org/10.1534/genetics.113.152462

Hoegh-Guldberg, O. (1999). Climate change, coral bleaching and the future of the world’s coral reefs. Marine and Freshwater Research, 50, 839–866.

Hoeksema, B. W. (2007). Delineation of the Indo-Malayan centre of maximum marine biodiversity: The Coral Triangle. In W. Renema (Ed.), Biogeography, Time, and Place: Distributions, Barriers, and Islands (pp. 117–178). Springer Netherlands. https://doi.org/10.1007/978-1-4020-6374-9_5

Horne, J. B. (2014). Thinking outside the barrier: neutral and adaptive divergence in Indo-Pacific coral reef faunas. Evolutionary Ecology, 28(6), 991–1002. https://doi.org/10.1007/s10682-014-9724-9

Jeffreys, H. (1998). The Theory of Probability. OUP Oxford. https://market.android.com/details?id=book-vh9Act9rtzQC

Johannesson, K., & André, C. (2006). Life on the margin: genetic isolation and diversity loss in a peripheral marine ecosystem, the Baltic Sea. Molecular Ecology, 15(8), 2013–2029. https://doi.org/10.1111/j.1365-294X.2006.02919.x

Kawecki, T. J. (2008). Adaptation to marginal habitats. Annual Review of Ecology, Evolution, and Systematics, 39(1), 321–342. https://doi.org/10.1146/annurev.ecolsys.38.091206.095622

Kelly, R. P., & Palumbi, S. R. (2010). Genetic structure among 50 species of the northeastern Pacific rocky intertidal community. PloS One, 5(1), e8594. https://doi.org/10.1371/journal.pone.0008594

Kirkendale, L. A., & Meyer, C. P. (2004). Phylogeography of the Patelloida profunda group (Gastropoda: Lottidae): diversification in a dispersal-driven marine system. Molecular Ecology, 13(9), 2749–2762. https://doi.org/10.1111/j.1365-294X.2004.02284.x

Kochzius, M., Seidel, C., Hauschild, J., Kirchhoff, S., Mester, P., Meyer-Wachsmuth, I., Nuryanto, A., & Timm, J. (2009). Genetic population structures of the blue starfish Linckia laevigata and its gastropod ectoparasite Thyca crystallina. Marine Ecology Progress Series, 396, 211–219. https://doi.org/10.3354/meps08281

Kool, J. T., Paris, C. B., Barber, P. H., & Cowen, R. K. (2011). Connectivity and the development of population genetic structure in Indo-West Pacific coral reef communities. Global Ecology and Biogeography: A Journal of Macroecology, 20(5), 695–706. https://onlinelibrary.wiley.com/doi/abs/10.1111/j.1466-8238.2010.00637.x

Kopelman, N. M., Mayzel, J., Jakobsson, M., Rosenberg, N. A., & Mayrose, I. (2015). Clumpak: a program for identifying clustering modes and packaging population structure inferences across K. Molecular Ecology Resources, 15(5), 1179–1191. https://doi.org/10.1111/1755-0998.12387

Kuhnt, W., Holbourn, A., Hall, R., Zuvela, M., & Käse, R. (2004). Neogene history of the Indonesian throughflow. Continent-Ocean Interactions within East Asian Marginal Seas. Geophysical Monograph, 149, 299–320. http://searg.rhul.ac.uk/pubs/kuhnt_etal_2004%20Neogene%20Indonesian%20throughflow.pdf

Leray, M., Beldade, R., Holbrook, S. J., Schmitt, R. J., Planes, S., & Bernardi, G. (2010). Allopatric divergence and speciation in coral reef fish: the three-spot dascyllus, Dascyllus trimaculatus, species complex. Evolution, 64(5), 1218–1230. https://doi.org/10.1111/j.1558-5646.2009.00917.x

Limborg, M. T., Helyar, S. J., De Bruyn, M., Taylor, M. I., Nielsen, E. E., Ogden, R., Carvalho, G. R., FPT Consortium, & Bekkevold, D. (2012). Environmental selection on transcriptome-derived SNPs in a high gene flow marine fish, the Atlantic herring (Clupea harengus). Molecular Ecology, 21(15), 3686–3703. https://doi.org/10.1111/j.1365-294X.2012.05639.x

Lin, T.-Y., & Liu, L.-L. (2008). Low levels of genetic differentiation among populations of the coral-inhabiting snail Coralliophila violacea (Gastropoda: Coralliophilidae) in regions of the Kuroshio and South China Sea. Zoological Studies, 47(1), 17. http://citeseerx.ist.psu.edu/viewdoc/download?doi=10.1.1.654.4420&rep=rep1&type=pdf

Lin, T.-Y., & Liu, P.-J. (1995). Fecundity of female coral-inhabiting snails, Coralliophila violacea (Gastropoda: Coralliophilidae). The Veliger, 38(4), 319–322.

Lischer, H. E. L., & Excoffier, L. (2012). PGDSpider: an automated data conversion tool for connecting population genetics and genomics programs. Bioinformatics, 28(2), 298–299. https://doi.org/10.1093/bioinformatics/btr642

Litton, C. D., & Jefferys, H. (1984). Theory of Probability (3rd Edition). Oxford University Press.

Liu, S.-Y. V., Chang, F.-T., Borsa, P., Chen, W.-J., & Dai, C.-F. (2014). Phylogeography of the humbug damselfish, Dascyllus aruanus (Linnaeus, 1758): Evidence of Indo-Pacific vicariance and genetic differentiation of peripheral populations. Biological Journal of the Linnean Society. Linnean Society of London, 113(4), 931–942. https://doi.org/10.1111/bij.12378

Liu, S. Y. V., Tuanmu, M. N., Rachmawati, R., Mahardika, G. N., & Barber, P. H. (2019) Integrating phylogeographic and ecological niche approaches to delimitating cryptic lineages in the blue-green damselfish (Chromis viridis). PeerJ., 7:e7384. doi:10.7717/peerj.7384

Longo, G., & Bernardi, G. (2015). The evolutionary history of the embiotocid surfperch radiation based on genome-wide RAD sequence data. Molecular Phylogenetics and Evolution, 88, 55–63. https://doi.org/10.1016/j.ympev.2015.03.027

Lourie, S. A., Green, D. M., & Vincent, A. C. J. (2005). Dispersal, habitat differences, and comparative phylogeography of Southeast Asian seahorses (Syngnathidae: Hippocampus). Molecular Ecology, 14(4), 1073–1094. https://doi.org/10.1111/j.1365-294X.2005.02464.x

Ludt, W. B., & Rocha, L. A. (2015). Shifting seas: the impacts of Pleistocene sea-level fluctuations on the evolution of tropical marine taxa. Journal of Biogeography, 42(1), 25–38. https://onlinelibrary.wiley.com/doi/abs/10.1111/jbi.12416

Martin, R., Hild, S., Walther, P., Ploss, K., Boland, W., & Tomaschko, K.-H. (2007). Granular chitin in the epidermis of nudibranch molluscs. The Biological Bulletin, 213(3), 307–315. https://doi.org/10.2307/25066648

Meyer, C. P., Geller, J. B., & Paulay, G. (2005). Fine scale endemism on coral reefs: archipelagic differentiation in turbinid gastropods. Evolution, 59(1), 113–125. https://www.ncbi.nlm.nih.gov/pubmed/15792232

Narum, S. R. (2006). Beyond Bonferroni: Less conservative analyses for conservation genetics. Conservation Genetics, 7(5), 783–787. https://doi.org/10.1007/s10592-005-9056-y

Nuryanto, A., & Kochzius, M. (2009). Highly restricted gene flow and deep evolutionary lineages in the giant clam Tridacna maxima. Coral Reefs, 28(3), 607–619. https://doi.org/10.1007/s00338-009-0483-y

Polato, N. R., Concepcion, G. T., Toonen, R. J., & Baums, I. B. (2010). Isolation by distance across the Hawaiian Archipelago in the reef-building coral Porites lobata. Molecular Ecology, 19(21), 4661–4677. https://doi.org/10.1111/j.1365-294X.2010.04836.x

Pritchard, J. K., Stephens, M., & Donnelly, P. (2000). Inference of population structure using multilocus genotype data. Genetics, 155(2), 945–959. https://www.ncbi.nlm.nih.gov/pubmed/10835412

Raynal, J. M., Crandall, E. D., Barber, P. H., Mahardika, G. N., Lagman, M. C., & Carpenter, K. E. (2014). Basin isolation and oceanographic features influencing lineage divergence in the humbug damselfish (Dascyllus aruanus) in the Coral Triangle. Bulletin of Marine Science, 90(1), 513–532. https://doi.org/10.5343/bms.2013.1017

Reid, D. G., Lal, K., Mackenzie-Dodds, J., Kaligis, F., Littlewood, D. T. J., & Williams, S. T. (2006). Comparative phylogeography and species boundaries in Echinolittorina snails in the central Indo-West Pacific. Journal of Biogeography, 33(6), 990–1006. https://doi.org/10.1111/j.1365-2699.2006.01469.x

Rocha, L. A., & Bowen, B. W. (2008). Speciation in coral-reef fishes. Journal of Fish Biology, 72(5), 1101–1121. https://onlinelibrary.wiley.com/doi/abs/10.1111/j.1095-8649.2007.01770.x

Saenz-Agudelo, P., Dibattista, J. D., Piatek, M. J., Gaither, M. R., Harrison, H. B., Nanninga, G. B., & Berumen, M. L. (2015). Seascape genetics along environmental gradients in the Arabian Peninsula: insights from ddRAD sequencing of anemonefishes. Molecular Ecology, 24(24), 6241–6255. https://doi.org/10.1111/mec.13471

Sanford, E., & Kelly, M. W. (2011). Local adaptation in marine invertebrates. Annual Review of Marine Science, 3, 509–535. https://doi.org/10.1146/annurev-marine-120709-142756

Sbrocco, E. J., & Barber, P. H. (2013). MARSPEC: ocean climate layers for marine spatial ecology: Ecological Archives E094-086. Ecology, 94(4), 979–979. https://esajournals.onlinelibrary.wiley.com/doi/abs/10.1890/12-1358.1

Shinoda, T., Han, W., Metzger, E. J., & Hurlburt, H. E. (2012). Seasonal variation of the Indonesian Throughflow in Makassar Strait. Journal of Physical Oceanography, 42(7), 1099–1123. https://doi.org/10.1175/JPO-D-11-0120.1

Simmonds, S. E., Chou, V., Cheng, S. H., & Rachmawati, R. (2018). Evidence of host-associated divergence from coral-eating snails (genus Coralliophila) in the Coral Triangle. Coral Reefs. 37(2), 355–371. https://link.springer.com/article/10.1007/s00338-018-1661-6

Simmonds, S. E., Fritts Penniman, A. L., Cheng, S. H., Mahardika, G. N., & Barber, P. H. (2020). Genomic signatures of host associated divergence and adaptation in a coral eating snail, Coralliophila violacea (Kiener, 1836). Ecology and Evolution, 10(4), 1817–1837. https://doi.org/10.1002/ece3.5977

Slatkin, M. (1993). Isolation by distance in equilibrium and non-equilibrium populations. Evolution, 47(1), 264–279. https://doi.org/10.1111/j.1558-5646.1993.tb01215.x

Soong, K., & Chen, J.-L. (1991). Population structure and sex-change in the coral-inhabiting snail Coralliophila violacea at Hsiao-Liuchiu, Taiwan. Marine Biology, 111(1), 81–86. https://doi.org/10.1007/BF01986349

Spalding, M. D., Fox, H. E., Allen, G. R., Davidson, N., Ferdaña, Z. A., Finlayson, M., Halpern, B. S., Jorge, M. A., Lombana, A., Lourie, S. A., Martin, K. D., McManus, E., Molnar, J., Recchia, C. A., & Robertson, J. (2007). Marine ecoregions of the world: A bioregionalization of coastal and shelf areas. Bioscience, 57(7), 573–583. https://doi.org/10.1641/B570707

Szab’o, Z., Snelgrove, B., Craig, M. T., Rocha, L. A., & Bowen, B. W. (2014). Phylogeography of the manybar goatfish, Parupeneus multifasciatus, reveals isolation of the Hawaiian Archipelago and a cryptic species in the Marquesas Islands. Bulletin of Marine Science, 90(1), 493–512. https://doi.org/10.5343/bms.2013.1032

Taylor, J. B. (1975). Planktonic prosobranch veligers of Kaneohe Bay, Hawaiian Island. Ph. D. Dissertation. Univ. Hawaii.

Teske, P. R., Sandoval-Castillo, J., Golla, T. R., Emami-Khoyi, A., Tine, M., von der Heyden, S., & Beheregaray, L. B. (2019). Thermal selection as a driver of marine ecological speciation. Proceedings of the Royal Society B: Biological Sciences, 286(1896), 20182023. https://doi.org/10.1098/rspb.2018.2023

Timm, J., & Kochzius, M. (2008). Geological history and oceanography of the Indo-Malay Archipelago shape the genetic population structure in the false clown anemonefish (Amphiprion ocellaris). Molecular Ecology, 17(18), 3999–4014. https://onlinelibrary.wiley.com/doi/abs/10.1111/j.1365-294X.2008.03881.x

Tornabene, L., Valdez, S., Erdmann, M., & Pezold, F. (2015). Support for a “Center of Origin” in the Coral Triangle: cryptic diversity, recent speciation, and local endemism in a diverse lineage of reef fishes (Gobiidae: Eviota). Molecular Phylogenetics and Evolution, 82 Pt A, 200–210. https://doi.org/10.1016/j.ympev.2014.09.012

Treml, E. A., Roberts, J., Halpin, P. N., Possingham, H. P., & Riginos, C. (2015). The emergent geography of biophysical dispersal barriers across the Indo-West Pacific. Diversity & Distributions, 21(4), 465–476. https://doi.org/10.1111/ddi.12307

Treml, E. A., Roberts, J. J., Chao, Y., Halpin, P. N., Possingham, H. P., & Riginos, C. (2012) Reproductive output and duration of the pelagic larval stage determine seascape-wide connectivity of marine populations. Integrative and Comparative Biology, 52(4) 525–537, https://doi.org/10.1093/icb/ics101

Voris, H. K. (2000). Maps of Pleistocene sea levels in Southeast Asia: shorelines, river systems and time durations. Journal of Biogeography, 27(5), 1153–1167. https://doi.org/10.1046/j.1365-2699.2000.00489.x

Waldrop, E., Hobbs, J.-P. A., Randall, J. E., DiBattista, J. D., Rocha, L. A., Kosaki, R. K., Berumen, M. L., & Bowen, B. W. (2016). Phylogeography, population structure and evolution of coral-eating butterflyfishes (Family Chaetodontidae, genus Chaetodon, subgenus Corallochaetodon). Journal of Biogeography, 43(6), 1116–1129. https://doi.org/10.1111/jbi.12680

Wang, S., Meyer, E., McKay, J. K., & Matz, M. V. (2012). 2b-RAD: a simple and flexible method for genome-wide genotyping. Nature Methods, 9(8), 808–810. https://doi.org/10.1038/nmeth.2023

Williams, S. T., & Benzie, J. A. H. (1998). Evidence of a biogeographic break between populations of a high dispersal starfish: Congruent regions within the Indo-West Pacific defined by color morphs, mtDNA, and allozyme data. Evolution, 52(1), 87–99. https://doi.org/10.1111/j.1558-5646.1998.tb05141.x

Wright, S. (1943). Isolation by distance. Genetics, 28(2), 114–138. https://www.ncbi.nlm.nih.gov/pubmed/17247074

